# Biofabrication of muscle fibers enhanced with plant viral nanoparticles using surface chaotic flows

**DOI:** 10.1101/2020.06.30.181214

**Authors:** Ada I. Frías-Sánchez, Diego A. Quevedo-Moreno, Mohamadmahdi Samandari, Jorge A. Tavares-Negrete, Víctor Hugo Sánchez-Rodríguez, Ivonne González-Gamboa, Fernando Ponz, Mario M. Alvarez, Grissel Trujillo-de Santiago

**Affiliations:** Centro de Biotecnología-FEMSA, Tecnológico de Monterrey, 64849 Monterrey, México; Mechatronics and Electrical Engineering Department, Tecnológico de Monterrey, 64849 Monterrey, México; Department of Biomedical Engineering, University of Connecticut, Farmington, CT 06030, USA; Bioengineering Department, Tecnológico de Monterrey, 64849 Monterrey, México; Centro de Biotecnología y Genómica de Plantas, Universidad Politécnica de Madrid - Instituto Nacional de Investigación y Tecnología Agraria y Alimentaria (CBGP, UPM-INIA), 28223 Madrid, Spain

**Author notes:** Corresponding authors. E-mail: GTdS MMA.

**Keywords:** bioprinting, hydrogel, biofabrication, tissue engineering, skeletal muscle, chaotic printing, viral nanoparticle

## Abstract

Multiple human tissues exhibit fibrous nature. Therefore, the fabrication of hydrogel filaments for tissue engineering is a trending topic. Current tissue models are made of materials that often require further enhancement for appropriate cell attachment, proliferation and differentiation. Here we present a simple strategy, based on the use surface chaotic flows amenable of mathematical modeling, to fabricate continuous, long and thin filaments of gelatin methacryloyl (GelMA).

The fabrication of these filaments is achieved by chaotic advection in a finely controlled and miniaturized version of the journal bearing (JB) system. A drop of GelMA pregel was injected on a higher-density viscous fluid (glycerin) and a chaotic flow is applied through an iterative process. The hydrogel drop is exponentially deformed and elongated to generate a fiber, which was then polymerized under UV-light exposure. Computational fluid dynamic (CFD) simulations are conducted to determine the characteristics of the flow and design the experimental conditions for fabrication of the fibers. GelMA fibers were effectively used as scaffolds for C2C12 myoblast cells. Experimental results demonstrate an accurate accordance with CFD simulations for the predicted length of the fibers.

Plant-based viral nanoparticles (i.e., *Turnip mosaic virus*; TuMV) were then integrated to the hydrogel fibers as a secondary nano-scaffold for cells for enhanced muscle tissue engineering. The addition of TuMV significantly increased the metabolic activity of the cell-seeded fibers (p*<0.05), strengthened cell attachment throughout the first 28 days, improved cell alignment, and promoted the generation of structures that resemble natural mammal muscle tissues.

Chaotic 2D-printing is proven to be a viable method for the fabrication of hydrogel fibers. The combined use of thin and long GelMA hydrogel fibers enhanced with flexuous virions offers a promising alternative for scaffolding of muscle cells and show potential to be used as cost-effective models for muscle tissue engineering purposes.

## 1. Introduction

Most organs in the human body are composed of fiber-like structures. Nerves, tendons, blood vessels, bones and muscles are some examples of tissues exhibiting a fibrous or tubular nature [1], which makes fibers a highly attractive architecture in the field of tissue engineering [2]. In particular, muscles make up around 40-45% of the lean body mass of an adult [3] and are essential for locomotion and posture [4,5], as well as for metabolic activity regulation [6]. Even though muscles are capable of regenerating themselves to some extent from mild damages [5], when serious injuries or intrinsic defects occur, additional strategies need to be implemented in order to improve muscle quality and maintain its functionality. To treat these conditions, some medical procedures, mainly autografts and heterologous implants, have been implemented. However, these surgeries are invasive, often produce complications in patients [7] and are greatly limited by tissue availability. Developing cost-effective artificial substitutes that adequately recapitulate the architecture and function of muscle tissues will enable the expansion of available solutions for tissue repair.

A wide variety of biofabrication methods for fiber production are available nowadays (i.e. 3D-bioprinting, electrospinning, microfluidics) [8–12]. Some of these methods have successfully been used for muscle tissue engineering [13–16]. Each of them has its advantages and limitations. For instance, 3D-bioprinting is a fast-developing technology, but the portfolio of biomaterials that possess the physicochemical properties required for the process are still limited [17], and are printed under a restricted resolution defined by each printer [18]. Electrospinning techniques can yield topographies that could induce enhanced cell adhesion and proliferation, but the commonly used materials have to be modified to provide them with cell-recognition signals [15], which are crucial for cells to attach, and the product is generally a dry polymeric structure that needs further materials to ensure cell survival. Microfluidic-based fabrication techniques can be used to generate long continuous filaments with various architectures, as well as multi-material structures [14]. However, the material selection is also limited and challenging, due to the fast solidification rate required for continuous fabrication [15]. Moreover, most of these techniques involve the use of extensive technical expertise and complex and usually expensive equipment. Although these technologies have reduced their costs throughout the years [19], the resolution obtained with the low-cost biofabrication methods to produce fibers is insufficient for building faithful 3D models of real tissues [20,21]. Furthermore, most state-of-the-art biofabrication systems involve complex multi-component equipment that may require high maintenance [21].

Tissue engineering offers a wide spectrum of alternatives for tissue repair, fabrication, and implantation. Among many possibilities, the interplay between biomaterials science and nanotechnology may contribute substantially to tissue engineering solutions [22,23]. Since cell behavior may be regulated by nanoscale features and several suitable materials for fiber fabrication lack essential components for cell attachment and proliferation, different nanotechnology approaches can be implemented in order to enhance these materials (i.e., nanoparticles, nanofibers, nanotopography). These strategies allow us to enhance the functioning of artificial tissue models by controlling cell-cell and cell-scaffold interactions [24]. However, the use of additional components can affect the biocompatibility and increase fabrication costs [25]. Plant-based viral nanoparticles (VNPs) are an attractive nano-tool for biomedical purposes since they are mainly made of protein, are harmless for mammals, can be extracted through processes that reach high yields, and can be produced relatively fast and at a low cost [26]. Some virions, like those of *Turnip mosaic virus* (TuMV) are especially attractive for tissue engineering purposes because of their flexibility and dimensions that are comparable to some components of the extracellular matrix [27,28].

In this work, a simple, efficient and cost-effective surface-printing method is demonstrated for the biofabrication of three-dimensional fibers for muscle tissue engineering purposes. With this aim, we used a miniaturized version of the Journal Bearing (miniJB) flow system, previously used to fabricate highly packed 3D-microstructure within small hydrogel constructs through a technique called chaotic 3D-printing [29]. In a novel embodiment of this technique, two liquids with different densities, namely glycerin and gelatin methacryloyl (GelMA), can be used to fabricate hydrogel fibers in surface 2D-flows under laminar conditions. The miniJB works as a microfabrication wheel to elongate the water-based pre-gel drop in an exponential fashion into a long and thin filament, which is then gelled by photopolymerization. Two versions of hydrogel fibers were assessed as scaffolds to fabricate our muscle fibers: (1) Fibers made of pristine GelMA inks, and (2) those fabricated using nano-composites made of GelMA/TuMV. TuMV nanoparticles were added to provide mechanical stability to the hydrogel fibers and to add a secondary nano-scaffolding structure that may serve as an amenable-to-functionalize platform to facilitate further increases of complexity. The performance of both types of fibers as scaffolds for muscle cells was tested.

## 2. Materials and methods

### Materials

Type A porcine skin gelatin, methacrylic anhydride (MA), Dulbecco’s phosphate-buffered saline (DPBS), 4′,6-diamidino-2-phenylindole (DAPI), paraformaldehyde, formaldehyde and penicillin-streptomycin (pen/strep) were purchased from Sigma-Aldrich. Phosphate-buffered saline (PBS), fetal bovine serum (FBS), and Dulbecco’s modified Eagle’s medium (DMEM) were purchased from Gibco. Lithium phenyl-2,4,6-trimethylbenzoylphosphinate (LAP) was purchased from Allevi, glycerin was purchased from a local supplier, murine C2C12 myoblasts were purchased from ATCC. Actin-Phalloidin iFluor 647 was purchased from AbCam, PrestoBlue® was purchased from Invitrogen, and ethanol (EtOH) was purchased from Hycel de México. CreateX fluorescent particles were purchased at an online store. TuMV virions were kindly provided by Fernando Ponz’s lab at Instituto Nacional de Investigación y Tecnología Agraria y Alimentaria (INIA), Madrid, Spain.

### GelMA synthesis

Type A porcine skin gelatin was mixed at 10% (w/v) in DPBS at 60°C. After being fully dissolved, 20% (w/v) of MA was incorporated dropwise in order to introduce the methacrylate substitution groups in the gelatin. The reaction was stopped by diluting four-fold with warm DPBS. In order to remove the residues of methacrylic acid, the mixture was continuously dialyzed for 5 days at 40°C using a custom-made dialysis system, later lyophilized for 5 days, and stored at −80°C until use.

### Ink preparation

All pregel inks were prepared with 10% (w/v) GelMA and 0.067% (w/v) LAP photoinitiator dissolved in DPBS at 60°C. Fluorescent particles were first washed by diluting 500 μl of fluorescent magenta CreateX particles in 14 ml of distilled water, followed by a thorough homogenization and a 5-minute centrifugation process at 1500 rpm. Excess water was replaced with new distilled water and the whole process was repeated twice. The remaining fluorescent particles were homogenized in 1 ml GelMA pregel. Virus-embedded inks included a 10 μg/ml concentration of TuMV virions, isolated from Indian mustard plants as described in the literature [30].

### MiniJB assembly

An inner mixing shaft of radius *r*_*in*_ and an external reservoir of radius *r*_*out*_ are the main cylindrical components that comprise the miniJB. The designed geometry of our system is based on two dimensionless parameters (Figure 1A): cylinder radii ratio, *r=r_in_/r_out_,* and the ratio of the eccentricity (*e*) to the outer cylinder radius, *ϵ=e/r_out_*. Two arrangements were used: the regular (*ϵ=0*) and the chaotic (*ϵ=1/3*), and for both *r=1/3*. The actual diameters of the external and internal cylinders were 15 mm and 5 mm, respectively. The process was carried out in cycles (Figure 1B), composed of two mixing actions: first, the counterclockwise rotation of the inner cylinder (⍺), followed by the clockwise rotation of the outer one (β). Of the many mixing protocols [⍺°, β°] available in the literature, we selected the [270°, 810°]. In order to adequately control the movement of each cylinder, we used an Arduino platform and 2 stepper motors. Our device was made of materials that can be easily found in hardware or online stores, and its overall cost is below 150 USD.

**Figure 1.**
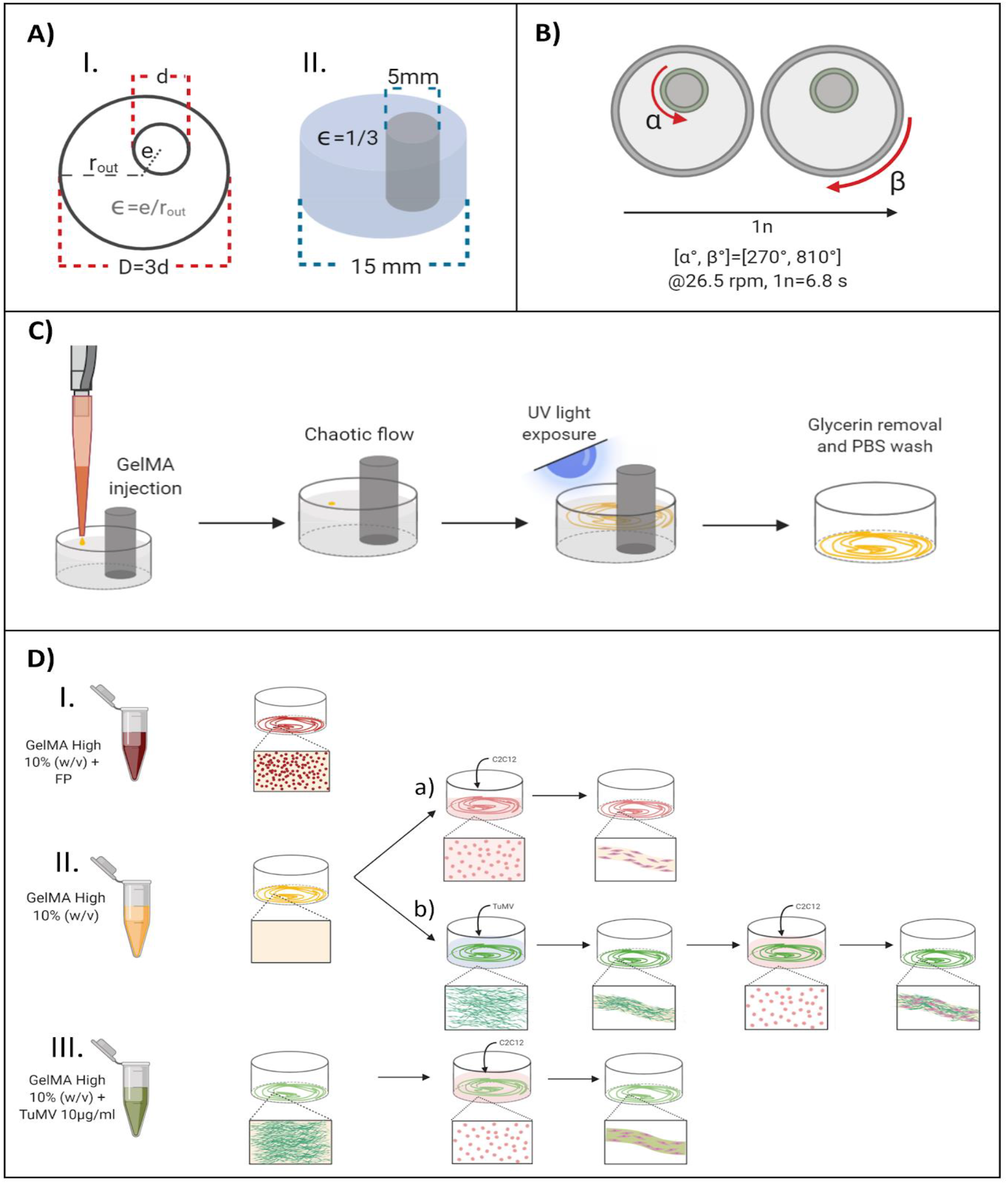
Experimental strategy. A) MiniJB device I) dimensionless parameters and II) actual dimensions, B) representation of one cycle of the process, C) schematic diagrams of fiber fabrication process and D) different types of fibers fabricated: I) GelMA with fluorescent particles, II) pristine GelMA a) cell-seeded or b) with a superficial TuMV coating followed by a cell-seeding process, and III) GelMA with embedded TuMV and subsequent cell-seeding.

### Fiber fabrication

We fabricated hydrogel fibers in a batch process using chaotic surface flows induced by the miniJB system. After the assembly of the device, 400 μl of glycerin (⍴=1.13 g/cm^3^) was placed inside the reservoir, and a 1 μl drop of GelMA pregel (⍴=1.01 g/cm^3^) was deposited on top. The device was later activated for a given amount of cycles, followed by a 15 s UV light exposure at 365 nm, glycerin drainage, and PBS was used to wash any fluid remainders (Figure 1C). In these experiments, the angular velocity for both cylinders, the mixing shaft and the reservoir of the system, was set to 26.5 rpm. All the process was carried out in sterile conditions inside a biosafety hood. The lengths of the fibers were measured with the Zen Blue Edition software (Carl Zeiss Microscopy GmbH, Germany) and plotted according to their cycle number. The collected data was used to determine the experimental *Lyapunov* exponent by calculating the slope of the linear portion from the plot of equation (1):

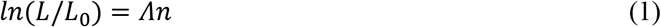

where *n* is the number of flow cycles, *L* is the length of the fiber after *n* cycles, and *L*_*0*_ is the original length of the filament (in this case, the diameter of the drop). All types of fibers used throughout the experimental process are shown in Figure 1D.

### TuMV coating: superficial virus

Hydrogel fibers, produced with a 1-cycle process, were placed in 24-well ultra-low attachment well plates and covered with a 10 μg/ml suspension of TuMV in PBS and incubated for 24 hours at 4°C prior to the cell-seeding process.

### Cell seeding

A 400 μl C2C12 cell suspension in DMEM growth medium (4×10^4^ cells ml^−1^) was seeded on all the chaotically-printed hydrogel fibers (1 cycle) in 24-well ultra-low attachment well plates. Fibers were incubated at 37°C in a humidified 5% CO_2_ atmosphere for 3 h to allow cell attachment on the fiber surface. The cell suspension was then replaced by DMEM growth medium with FBS 10% (v/v). Metabolic activity, as well as the cell morphology and alignment were monitored at different days.

### CFD simulations

Numerical modelling was implemented to determine relevant characteristics of the chaotic flow. First, the Navier-Stokes equations, formulated for incompressible laminar flows in a rotating coordinate system, were solved in the well-known journal bearing (JB) geometry. Two rotating domains were used to describe the independent rotation of the internal and external cylinders. The dynamics on the transition time-points were neglected in our simulations, meaning that we assumed that convection at each cycle ceased immediately after stopping each mixing action (i.e., inertial effects after each cycle were neglected). Simulations were performed in COMSOL Multiphysics® 5.4 (Comsol Inc., MA, USA). Two velocity fields were solved for inner and outer rotations using “Rotating Machinery, Laminar Flow” physics and “Frozen Rotor” stationary solvers. A fluid viscosity of 1 Pa.s and density of 1.12 g ml^−1^ were considered for the material properties of glycerin at room temperature. In each rotation, a non-slip boundary condition was considered for the fixed walls, and a rotating wall condition was applied to the wall of the moving cylinder. Subsequently, a particle tracking algorithm with massless formulation was implemented in the pre-calculated velocity fields to evaluate the dynamics and the position of all points composing the GelMA solution interface. A freeze boundary condition was considered in particle tracking and a time-dependent solver was used with multiple steps corresponding to multiple rotations to track the position of up to 50,000 particles in the system. For the estimation of the *Lyapunov* exponent in our 2D-chaotic flow protocols, we simulated the deformation of 1 mm lines (i.e., collections of points) located at different radial positions within the JB surface and calculated the value as we did with the fabricated fibers.

### Metabolic activity analysis

Cell viability was indirectly assessed by analyzing cell metabolic activity on days 0, 1, 2, 3, 6, 9, 12, 18, and 24 using the PrestoBlue^®^ assay. Briefly, fibers were covered in DMEM culture medium with 10% v/v PrestoBlue^®^ reagent and incubated for 2 h at 37°C. In a 96-well plate, 100 μl of medium were dispensed, and the fluorescence was measured in a microplate reader (Synergy™ HTX Multi-Mode Microplate Reader, BioTek Winooski, VT, USA) at 530/570 nm excitation and emission wavelengths. Fluorescence readings were normalized with respect to a control well with medium in the absence of samples.

### Staining of actin filaments and cell nuclei

The cell morphology was analyzed using fluorescent staining methods on days 1, 7, 14, 21, and 28. Briefly, the cell-laden fiber was fully covered with 4% paraformaldehyde solution in PBS for 30 min at room temperature to fix the cells. Samples were later stained with 1X Phalloidin iFluor 647 for filamentous F-actin, and 1 μg/ml DAPI-PBS for cell nuclei. After 90 min of incubation at room temperature, the samples were examined in an inverted microscope (Axio Observer Z1, Carl Zeiss Microscopy GmbH, Germany), using the Zen Blue edition software.

### Biofabricated fiber sample preparation (SEM)

The microarchitecture of our samples was analyzed on days 1 and 14 by SEM. The sample preparation involved a cell-fixation process and a dehydration process. The former consisted of a 4% paraformaldehyde followed by a 4% formaldehyde sample submersion, and the latter by a 4-hour ethanol-based graded dehydration process. Once samples were fully dry, they were sputtered with a 5 nm gold layer using a Q150R ES rotating sputter machine (Quorum, United Kingdom) to make the surface electrically conductive. These samples were analyzed using an EVO^®^ MA 25 SEM (Carl Zeiss Microscopy GmbH, Germany).

### Real muscle tissue sample preparation (SEM)

Calf gluteobiceps and gracilis muscles were dissected and submerged in PBS with 1% pen/strep, cut in 1 cm^3^ cubes and gradually frozen until reaching −80 °C. The fixation process for SEM analysis of these samples was the same as the previously described. Real tissue samples were dehydrated for 8 hours and gold-coated with a layer of 10 nm thickness.

### Image analysis

The number of nuclei and the cell orientation were determined from fluorescent microscopy images using Fiji/ImageJ (National Institutes of Health, AZ, USA) software. In order to determine the nuclei density, 15 sections of 100×100 μm were analyzed per micrograph. For the orientation analysis, cells whose direction was within ±10° with respect to the fiber longitudinal axis, were considered aligned. Angles were normalized to 0° and plotted in polar graphs using a custom-made program in MATLAB (MathWorks, MA, USA).

### Statistical analysis

All the experimental data regarding fiber dimensions was reported as mean ± standard deviation (SD). The statistical analysis for the metabolic activity assay data was performed using one-way ANOVA, and the nuclei density statistical analysis was assessed with two-way ANOVA, with significance levels of (*) p<0.05, (**) for p<0.01, and (***) for p<0.001.

## 3. Results and discussion

### Fiber fabrication

Long GelMA fibers were fabricated using surface 2D flows generated by a lab-made miniJB system as described in Figure 1. The JB flow was first described by Swanson and Ottino [31] as a simple flow system capable of producing chaotic flows. Here we have miniaturized this system to enable the creation of hydrogel fibers of thicknesses in the range of hundreds to tens of micrometers. To that aim, we generated a 3D flow with a viscous fluid (glycerin), on top of which we placed a small GelMA pregel drop of approximately 2.5 mm in diameter. The difference in density between both fluids is key for this method, since it allows the pregel drop to remain on the surface of the glycerin during the whole process and enables a 2D deformation of the hydrogel. In this work we used two miniJB setups: the concentric and the eccentric. When the two cylinders are concentric, the flows generated are regular and caused a linear elongation, while the eccentric arrangement generates chaotic flows that cause an exponential stretching.

Representative fluorescence micrographs in Figure 2A display the resulting fibers achieved through the regular and chaotic setups for the first two cycles. Evidently, the fibers fabricated using regular flows were substantially shorter than those produced by chaotic flows at the second cycle. Additionally, the thicknesses observed in linearly stretched fibers also display more consistent widths along the structure. In this particular approach, the glycerin is used exclusively as a vehicle for elongation of the GelMA drop. Glycerin subsequent drainage, as well as the removal of the mixing shaft, causes the fiber to modify the shape achieved at the end of the printing process. However, the dimensions of the fibers were accurately described by our CFD simulations.

**Figure 2.**
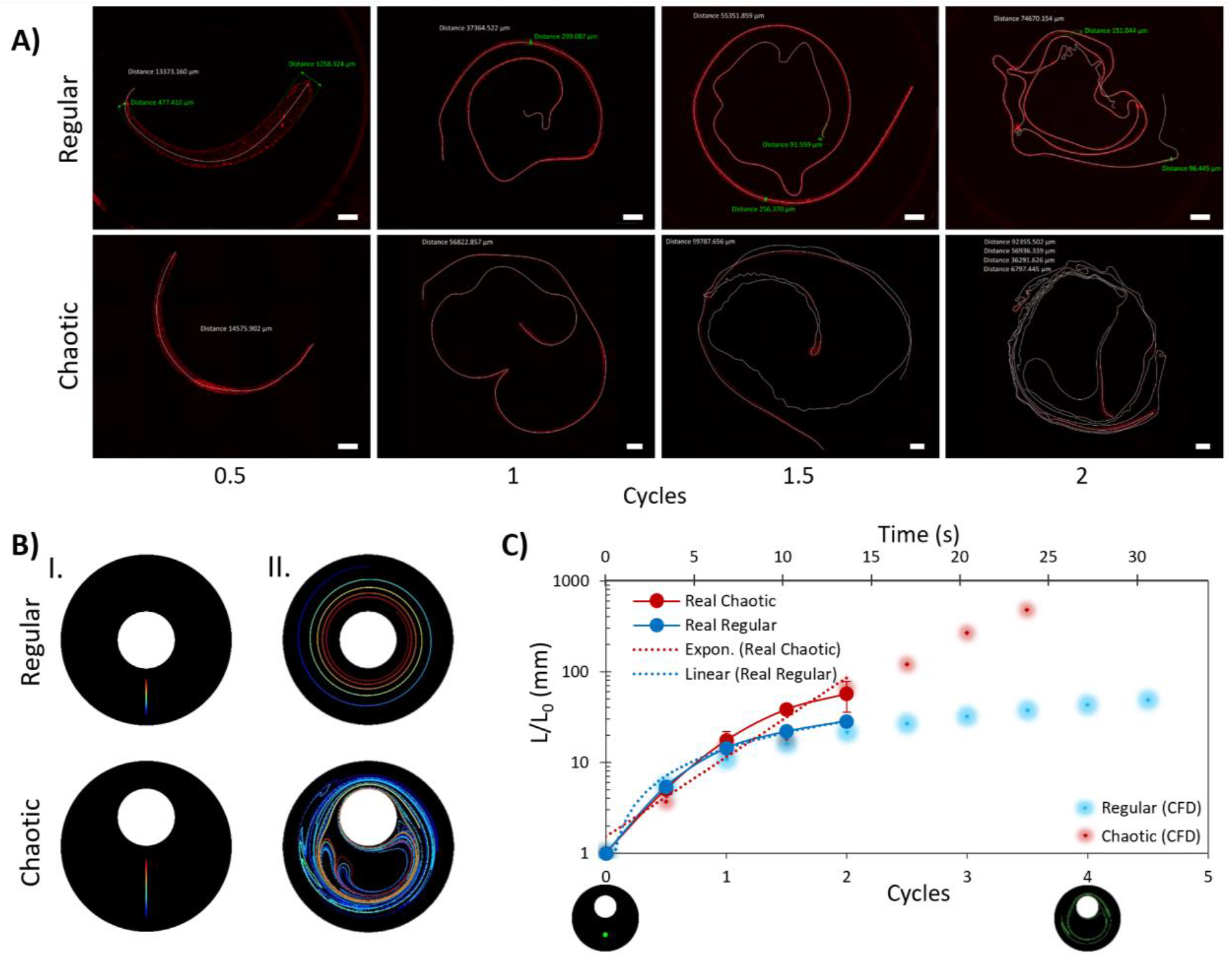
Chaotically printed fibers and CFD simulations. A) Fluorescence micrographs of GelMA fibers fabricated through regular and chaotic arrangements and at different numbers of cycles in a miniJB device [270°, 810°]. Scale bar: 1000 μm, B) CFD simulations of the deformation of centerlines caused by regular and chaotic flows I) before and II) after 5 cycles [270°, 810°], C) comparison between elongation of experimental fibers against CFD predictions.

Table S1 shows the lengths of fibers fabricated using different numbers of cycles in regular and chaotic flows generated by our miniJB system. High standard deviations can be attributed to initial differences in the drop placement, accidental losses of fiber portions due to experimental manipulation (these hydrogel fibers are extremely fragile), and intrinsic limitations of the optical microscopy equipment. Results reveal that our single initial drops were turned into filaments in average 56-fold longer after only two cycles of exposure to chaotic flow. In contrast, filaments stretched through the regular process only increased their length by a 28-fold in average (Table S2).

Conventional 3D-bioprinters generally generate structures at a linear rate, while our chaotic 2D-printer does it exponentially fast and at a noticeably lower cost. Therefore, chaotic 2D-printing can fabricate fibers of a given length much faster than a 3D bioprinter. Interestingly, the time economy achieved by chaotic printing is more evident as the number of cycles is increased, since the process of elongation of the hydrogel filament is exponential in chaotic flows. Note that the volume of the initial drop of GelMA is preserved during the process of elongation. Therefore, as the fiber elongates, its thickness is reduced. After a few cycles, we can potentially achieve fiber thicknesses in the nanoscale range [29]; however, handling hydrogel fibers of these dimensions is a challenging task. Furthermore, the resolution of the currently employed methods for the analysis of the length and width of our samples will eventually be insufficient to document remarkably thin filaments, since this fabrication technique induces an exponential contraction in thickness as the number of cycles is increased.

Relevant fiber dimensions for skeletal muscle tissue engineering can be achieved since the first cycle. Chaotic 2D-printing can produce fibers with varying diameters (Figure S2), as a result of the different levels of stretching that specific sections of the fiber undergo. These variations can be beneficial for the development of muscle-fiber models that can be used for specific tissue engineering applications. For instance, the depressor anguli oris, a facial muscle, exhibits a maximum length of 48 mm and width ranging from 8.8 to 35 mm [32]. This muscle could be built with chaotically printed fibers in a bottom-up approach. Fibers with even diameters can also be useful and can easily be achieved through the regular arrangement in the same miniJB system. Since we can design appropriate protocols to achieve specifically aimed dimensions, we can adapt our printing process for the fabrication of fibers with different dimensions.

### Computational fluid dynamics (CFD) simulations

Deterministic flows, whose dynamics are governed by the Navier-Stokes equations of motion, were used for the chaotic printing process. Therefore, the deformation and elongation caused by the chaotic flow can be mathematically modeled. The geometry of the system, the physical properties of the materials used, and the low-speed angular velocity yield a laminar surface flow that can be modeled as a 2D flow, which allows a numerical calculation of the velocity fields using CFD. Chaotic flows are capable of generating aligned microstructure at an exponential rate [33,34] due to their inherent properties, namely, the existence of a unique flow intrinsic skeleton, asymptotic directionality, and exponential divergence of points initially adjacent.

Multicolored lines (i.e., collections of particles) shown in Figure 2B were tracked both for the regular and chaotic simulations of the miniJB system flows throughout the first 5 cycles. This simple exercise, with relatively low computing investment, provided an accurate prediction of the structure observed at the top layer of the working fluid and enabled the estimation of the *Lyapunov* exponent (Λ) of the surface chaotic flow. A profound physical meaning is conveyed through the Λ value; this parameter describes the overall potential (and rate) of a chaotic mixing protocol to produce elongation and generate structure. CFD simulations determined a Λ value of 1.97 cycle^−1^ for the first two cycles, which closely matched the one calculated from the experimental data (2.01 cycle^−1^). In order to make a direct comparison, the data gathered from the CFD simulations and the experiments regarding the elongation of the drops are shown in Table S2. The experimental length has been normalized with respect to the diameter of the initial drop located in the system. The correlation between the experimental and simulated data was R^2^=0.9289, which indicates a high level of accuracy in the CFD predictions. A direct comparison of the simulated and experimental results is presented in Figure 2C.

Altogether, results demonstrate that we can reproduce the behavior of the actual experimental system using CFD simulations; therefore, our simulations enable the prediction (and design) of the filaments to be produced. However, these simulations also highlighted low movement areas, which indicate that the initial drop position can induce some level of variations in the length of the final fiber, particularly in initial cycles. In order to determine the level of stretching that each part of the centerline undergoes throughout the printing process, a material line composed of 50,000 particles was tracked throughout four cycles. Figure S4 shows the distances between these particles at an initial stage and after 2 and 4 cycles. Particles located between particles 1 and 25,000 exhibit relevant increases in the distance between neighboring particles both after the second and fourth cycles in most sections, in comparison to their original position. However, the distance between the particles located in the 25,000 - 34,000 range did not change significantly after two cycles and some (i.e., 27,000-28,000) even get closer to each other after the fourth cycle. Therefore, the initial drop location can be another parameter for controlling the length of the final fiber in the proposed system.

### Hydrogel fibers as cell scaffolds

Once a robust fabrication process for GelMA fibers was established, we tested different fabrication strategies for developing fibers containing mammalian cells (Figure 3A). We adopted C2C12 myoblasts as our cellular model. A first strategy, possibly the most intuitive, consisted in formulating a cell-laden bioink (i.e., suspending a cellular pellet in 10% GelMA) and depositing a drop of this bioink as an initial condition for our chaotic elongation process. However, preliminary assays with cell-laden hydrogel drops showed low post printing cell viability, possibly due to the shear stress that cells undergo during the elongating process. In any location of a chaotic flow, the stretching intensity can vary by several orders of magnitude, and C2C12 cells are sensitive to these conditions [35]. Consequently, we opted to use the produced hydrogel as cell-scaffolds and seeding C2C12 cells on the surface of the GelMA fibers soon after fabrication (Figure 1D ii and iii). We selected TuMV nanoparticles because their protein-based nature and their dimensions and flexibility could function as morphological cues that resemble some fibrillar components of the extracellular matrix, such as elastin and proteoglycans [27,28]. Three variants of this strategy were investigated. In the first one, cells were surface seeded on pristine 10% GelMA. In the second, GelMA was surface treated with TuMV nanoparticles. In a third variation, TuMV virions were suspended in the hydrogel prior to the printing process.

**Figure 3.**
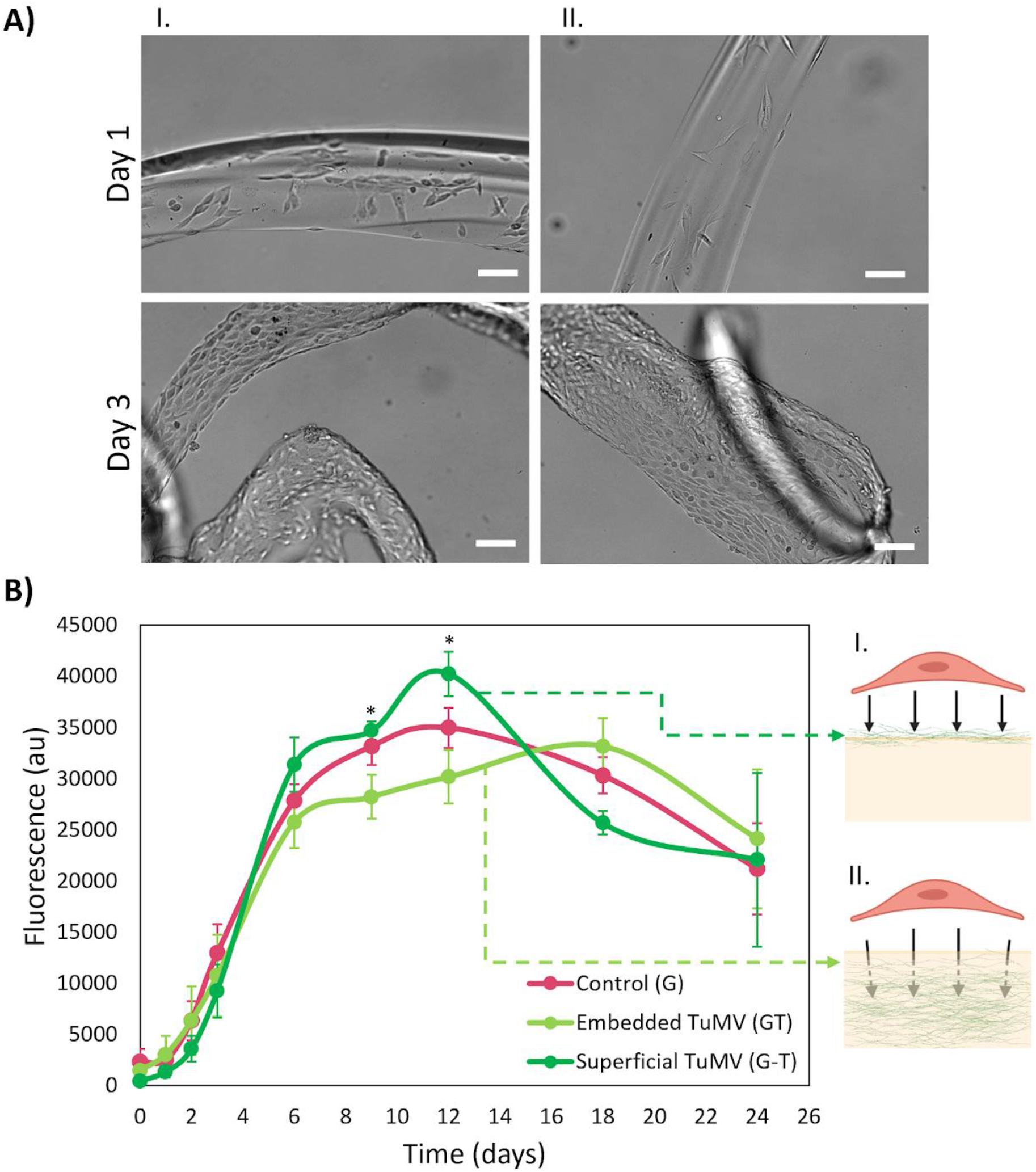
Cell-seeded fiber samples. A) Brightfield micrographs of C2C12 cells seeded on I) pristine GelMA fibers and II) GelMA fibers with superficial TuMV at days 1 and 3. Scale bar: 100 μm. B) Fluorescence readings from PrestoBlue^®^ assay (p*<0.05) with schematic representation of C2C12 cells interacting with I) superficial and II) embedded TuMV.

Figure 3B shows the PrestoBlue^®^ assay results, conducted to measure the metabolic activity of our cell-seeded fibers. We observed a gradual and substantial increase in activity over the first 12 days in all cases (i.e., GelMA fibers with or without TuMV), which is expected at proliferation stages and has also been reported for C2C12 cells seeded on GelMA scaffolds [36,37]. All samples followed approximately the same trend the first 6 days, but in the following days they exhibited significantly different behaviors (p*<0.05). Fibers with superficial TuMV displayed a higher metabolic activity on days 9 and 12, which could be the result of interactions between the cells and the readily exposed viral nanoparticles. On the other hand, fibers with embedded TuMV had a steadier and slower metabolic activity increase, with a peak of activity on day 18, unlike the other samples, which had an earlier metabolic activity peak on day 12. This could possibly be explained by the hypothesis of cells taking longer to be in contact with the TuMV nanoparticles in the embedded setup.

Fluorescence microscopy images in Figure 4 show strong evidence of the successful attachment, migration and proliferation of C2C12 cells on both pristine GelMA and GelMA/TuMV scaffolds. TuMV nanoparticles seem to produce an enhancing effect on the attachment of the cells to the hydrogel. On day 1, cells seeded on pristine GelMA fibers appear to display a higher organization than the ones on nanoenhanced scaffolds. Cell migration depends both on inner contractile forces and the adhesive force between cells and the scaffolds [38]. Stronger cell attachment would result in slower migration velocities, which would explain the initial difference between the morphology and arrangement of cells in both types of fibers. The hypothesis of improved attachment derived from the TuMV virions is further supported by the superior stability of the multinucleated cells on days 14 through 28. In this period, fibers containing TuMV at their surface retained a greater number of cells attached than the fibers fabricated with pristine GelMA. This superior performance was especially noticeable on day 28, where cells visibly detached from pristine GelMA fibers. Furthermore, as the myoblasts started elongating and fusing with each other, the resulting multinucleated cells self-aligned with respect to the longitudinal axis of the fiber. This spontaneous alignment agrees with findings from Costantini *et al.* where a relationship between the level of confinement and the orientation of C2C12 cells is reported [39]. At day 21, we observed highly dense tissue constructs that exhibit high degrees of cell alignment in both types of filaments. Even though many cells failed to remain adhered to the scaffolds made with pristine GelMA throughout the whole experimental period, the cells that did stay attached to the hydrogel kept their orientation.

**Figure 4.**
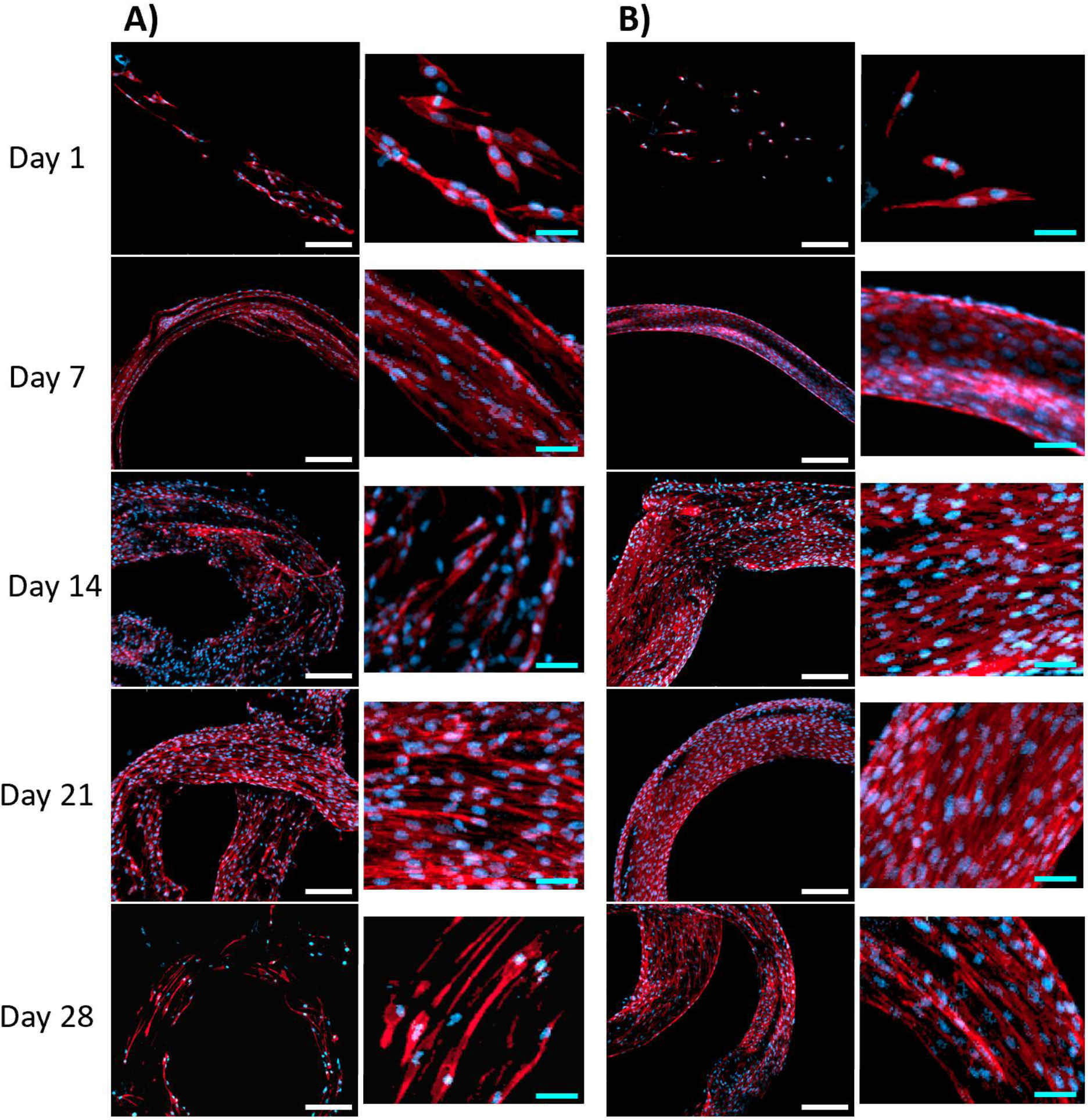
Fluorescence stainings of C2C12 cells cultured on A) GelMA and B) GelMA fibers containing superficial TuMV. F-actin shown in red and nuclei in blue. White scale bars: 200 μm, blue scale bars: 50 μm.

### Image analysis

An increase in nuclei density is one obvious indicator of cell population growth that can be easily measured through image analysis techniques. We analyzed the fluorescence micrographs using Fiji/ImageJ (open source software) and found that the nuclei density was initially increasing rapidly (days 1 to 7). Previous studies have reported the importance of the stiffness of scaffolds for an appropriate development of C2C12 cells. Gatazzo *et al.* used gelatin-based hydrogels and concluded that reproducing a native-resembling stiffness of scaffolds for muscle tissue engineering increases proliferation and differentiation of C2C12 cells [40], and reported an equivalent increase in cell nuclei to the one shown in Figure 5A. Our working concentration of GelMA (10% w/v) usually exhibits a stiffness of ~15 kPa [41], which is also similar to the native conditions of muscles (10-15 kPa [41]). Moreover, these results support the previously established hypothesis of an enhancement in cell attachment produced by the superficial TuMV during the last stage of the experiments (days 21 and 28), by displaying an apparent stability on TuMV-containing fibers, which is expected for a consistent cell population. A drastic decrease in nuclei density is observed in the pristine-GelMA fibers over the same period, where samples displayed a significantly lower nuclei density than their nanoenhanced counterparts (p***<0.001 on day 28). Interestingly, the cell nuclei density graphs are consistent with the trend observed throughout the whole duration of the PrestoBlue^®^ assay. The initial cell population growth explains the fast metabolic activity increase during the first 14 post-seeding days and the later decrease in cell density (days 21 to 28) is probably the cause of the final reduction in metabolic activity.

**Figure 5.**
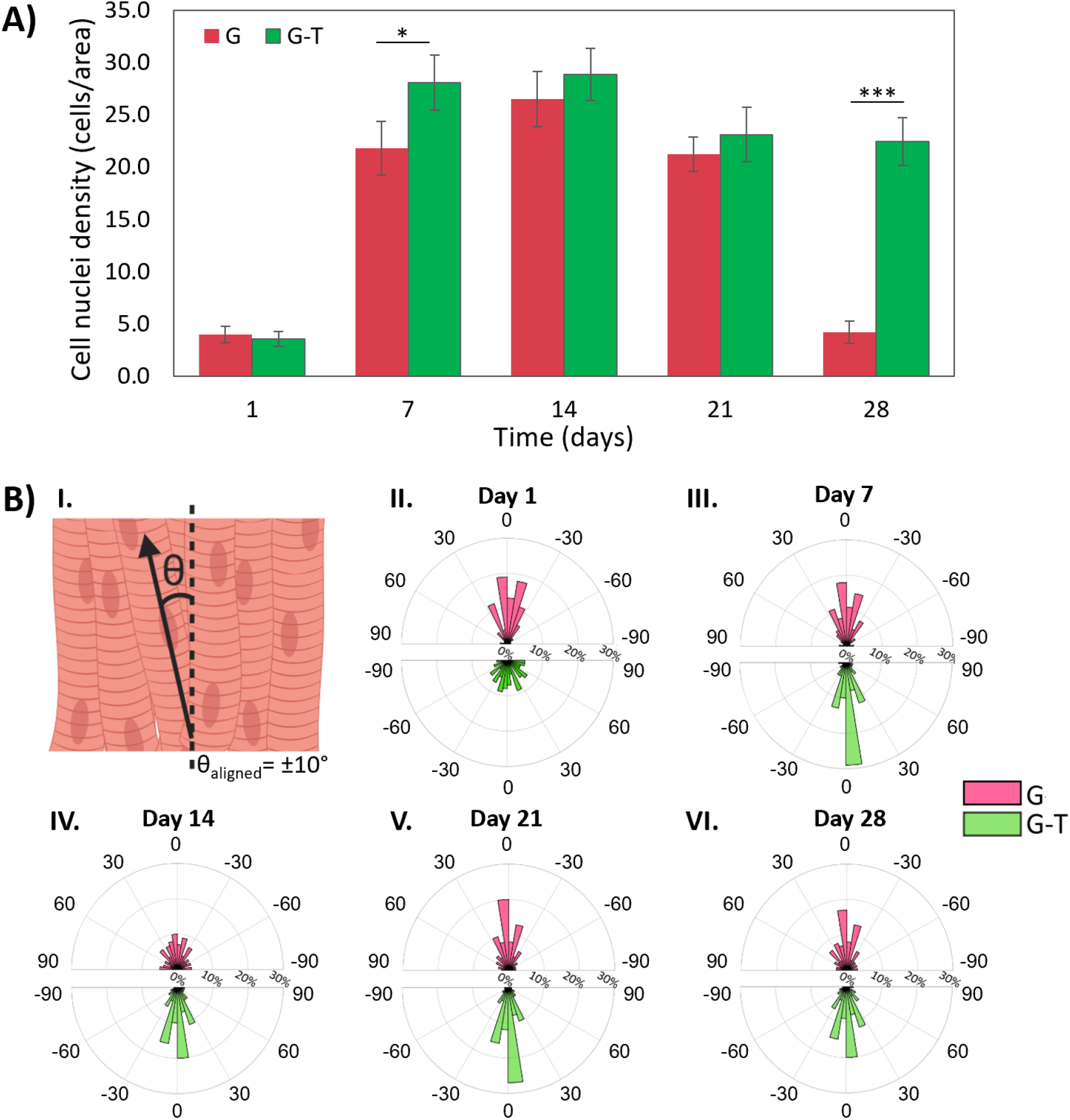
A) Cell nuclei density of GelMA and GelMA-TuMV fibers throughout time. P-values p*<0.05, p***<0.001, B) Cell orientation analysis I) Schematic representation of axial reference used for the determination of angles, and II-VI) polar graphs comparing cell-orientation for C2C12 cells cultured on GelMA and GelMA-TuMV fibers.

Cell alignment is one of the key characteristics that need to be achieved in order to have an artificial muscle that has potential to be functional, since it enables appropriate force transmission and contractile responses to the electrical stimuli provided by motor neurons [42]. The geometrical confinement where myocytes are enclosed, as well as the topography of the substrates where cells are seeded, have been proven to be relevant in their resulting orientation. Several techniques have been implemented in hydrogel scaffolds to improve cell orientation. Ebrahimi *et al.* fabricated micropatterned GelMA fibers (30% w/v) of ~500 μm through microfluidic spinning and demonstrated that the topographical cues, coupled with an agrin (a proteoglycan) treatment, have a beneficial and synergistic effect in cell orientation, backed up by an outstanding ~90% alignment of cells seeded superficially [43].

In another recent study, Chen *et al.* provided mechanical stimulation to C2C12 cell-laden GelMA fibers (10% w/v) and reported that ~80% of cells aligned within ±10° [44]. Various studies have found that wider structures hinder C2C12 myotube orientation [39,45–47], and plain hydrogels have been widely used as control groups to show that the lack of micro- and nanopatterns or bounding barriers in large scaffolds results in very low degrees of alignment (~10%) [47–50]. In our study, the nanoenhanced GelMA fibers were seemingly randomly oriented at the beginning, as opposed to the GelMA ones that showed a relatively higher organization (Figure 5B). However, a progressively increasing degree of alignment was henceforth observed on our fiber samples, both for GelMA and GelMA-TuMV. Our orientation analysis indicates that after day 7, in average, ~50% of cells located in nanoenhanced samples successfully aligned, while only ~40% of the cells in pristine-hydrogel fibers did. Both types of fibers showed considerably higher degrees of alignment than the aforementioned control groups from the literature. Remarkably, this was accomplished in the absence of any sort of external stimulation. We can thus suggest that the elongated and narrow architecture of our fibers induces cell alignment.

Cooperative approaches have also been performed by combining diverse sets of stimuli (i.e., topological, thermal, electrical, or mechanical) to enhance cell alignment [44,45,47,50–53]. Ahadian *et al.* improved myotube orientation from ~50% to ~80% by electrically stimulating micropatterned hydrogel samples with an interdigitated array of electrodes. Therefore, we anticipate that by providing further stimulation to our fibers, we can achieve higher degrees of cell alignment. All in all, our experimental data confirms that C2C12 cells attach to the fiber-scaffold, successfully spread, proliferate, and align over time.

### Scanning electron microscopy

In order to study the topography and internal microstructure of our fibers, we conducted SEM analysis. Figure 6 presents superficial and cross-sectional SEM micrographs from the chaotically printed fibers at days 1 and 14 after seeding. These images reveal that initially, fibers are sparsely populated and do not exhibit relevant cell organization. GelMA-TuMV fibers showed higher numbers of cells on their surface in comparison with the pristine GelMA samples. This difference further supports the hypothesis of an attachment-enhancing effect of the virions on the hydrogel, since cells on the pristine GelMA fibers would detach more easily from the scaffolds during the preparation process for the SEM analysis. On day 14, striations can be seen on the surface of both types of fibers, which are similar to those observed in the musculoskeletal hydrogel constructs fabricated by Heher and colleagues in a bioreactor that provides mechanical stimulation [50]. Cross-sectional views of our samples allow visualization of inner granular structures of similar dimensions to dehydrated myotubes from real muscle, which could suggest cell penetration into the hydrogel scaffold.

**Figure 6.**
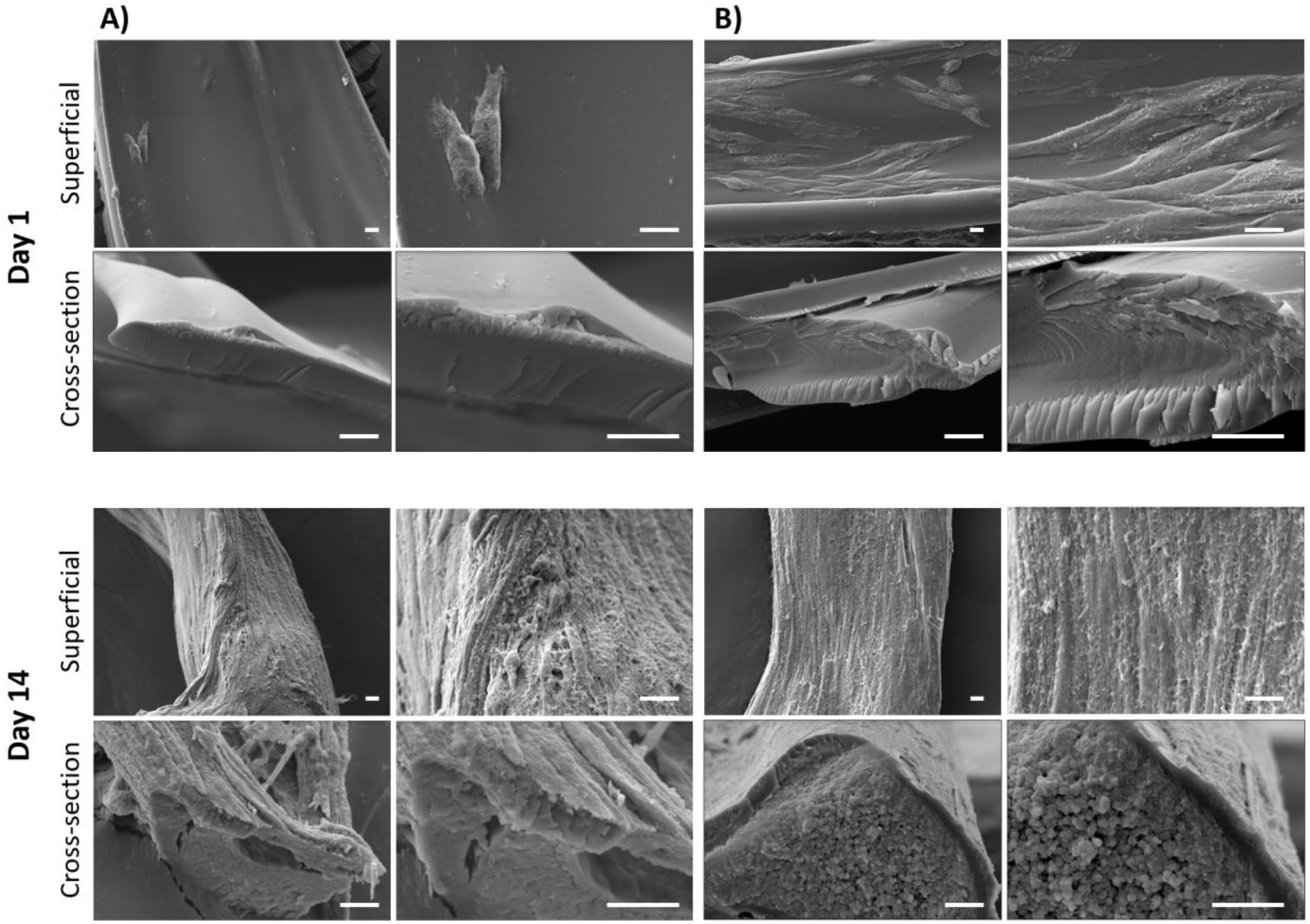
Superficial and cross-sectional SEM characterization. Cross-sectional and superficial SEM micrographs of A) GelMA and B) GelMA-TuMV fibers. Scale bars: 10 μm.

Several factors in the microenvironment affect the morphology of tissues. In Figure 7, a straightforward comparison between our nanoenhanced muscle tissue-engineered fibers and a real mammal muscle tissue is made. Characteristic fibrillar-like architectures observed in the native tissues can also be found in our lab-made fibers within similar scales, which suggests a structural and biological relevance. Earlier studies based on decellularized mammalian tissues showed samples that exhibit these threadlike structures. Wolf *et al.* worked with decellularized tissues to test the differences between specific and non-specific scaffolds for skeletal muscle repair [54]. DeQuach *et al.* used a similar approach for the development of injectable hydrogels for the treatment of muscle ischemia [55]. Both studies present SEM micrographs of the scaffolds based on decellularized tissues that highlight this structural composition based on tubular structures. These samples showed a great regenerative performance when tested *in vivo*. The current resemblance of our lab-made muscle and the native counterpart, just like the cell alignment, can arguably be improved by providing further stimulation (i.e., mechanical, electrical), in order for the tissue to mature in a more physiologically relevant manner.

**Figure 7.**
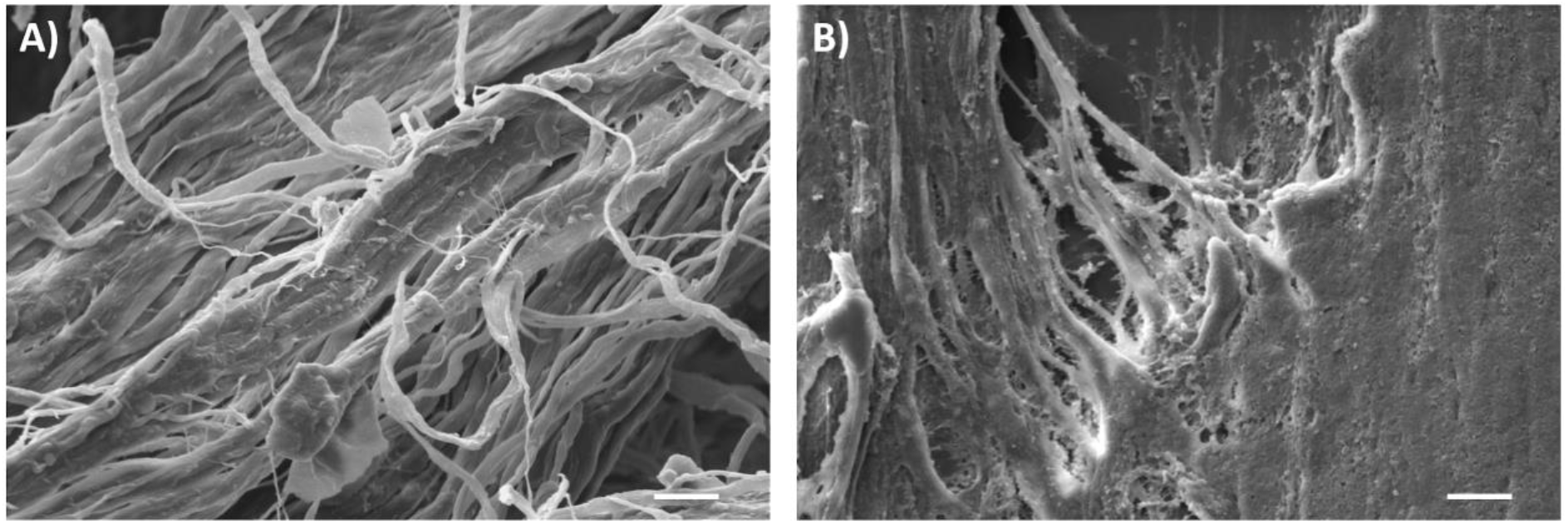
Comparison between real and artificial muscle tissue. SEM micrographs from A) native mammal muscle tissue and B) lab-made fiber (GelMA-TuMV) after 14 days of culture. Scale bar 10 μm.

## 4. Conclusions

We demonstrated that our miniJB is a low-cost system that can effectively produce predictable 2D surface chaotic flows and exploit their intrinsic potential to fabricate fibers exponentially fast by dispensing a lower density ink onto the surface of a higher density fluid contained in a reservoir. Our printing protocols are rooted in strong fundamental concepts of chaotic mixing and since their dynamics are governed by numerically solvable equations of motion, we can predict our outcomes by conducting CFD simulations. Our experimental and computational results led to closely matching *Lyapunov* values for the first 2 cycles (2.01 and 1.98, respectively) and produced a correlation coefficient of R^2^=0.9289. However, the manipulation of the fibers produced, as well as the techniques used to analyze their dimensions, will become a challenge as the number of cycles increases and the fiber diameter narrows down to the submicron scale.

Our technological platform can be used for tissue engineering applications by using hydrogels (i.e., GelMA) as inks and seeding cells on the formed scaffolds. We produced GelMA fibers and seeded musculoskeletal C2C12 murine myoblast cells on their surface. The chaotically printed fibers used in our experiments were proven to be effective scaffolds for these cells. The cells were metabolically active after being seeded on the scaffolds and increased their activity throughout the first 12 days. Cells were able to proliferate and fully populate the fibers, fused to presumably form myotubes, and oriented themselves to form aligned structures that kept their orientation over time.

The effect of TuMV was beneficial in terms of cell adhesion, alignment and long-term survival. Nanoenhanced fibers showed a significantly higher metabolic activity on day 12 (p*<0.05) and demonstrated greater stability by preserving their structure for at least 28 days, while pristine-GelMA fibers caused some cells to detach from the scaffolds after day 14. Cell orientation averaged ~50% after day 7 in fibers containing TuMV, which is ~10% above the percentage accomplished by the GelMA ones. Finally, our nanoenhanced chaotically printed muscle fibers displayed fibrillar-like architectures similar to real animal muscle tissue, which suggests structural and biological relevance of our models.

## Author contributions and acknowledgements

### Author contributions

GTdS and MMA conceptualized the research. GTdS, MMA, AIFS and DAQM designed the experiments. IGG extracted the TuMV virions and provided expertise for the experimental design. FP provided essential reagents and technical resources. VHSR produced the hydrogel for the experiments. MS performed the CFD simulations. AIFS and DAQM, executed the fabrication and cell culture experiments. JATN collaborated in cell culture and SEM experiments. AIFS, GTdS, and MMA analyzed the data, produced the figures, and wrote the manuscript. All authors reviewed and edited the manuscript before submission.

## Acknowledgements

This work was supported by Consejo Nacional de Ciencia y Tecnología (CONACyT) and Tecnológico de Monterrey. FP thanks the Spanish Ministry of Science for the Severo Ochoa Excellence Accreditation to the Centre for Plant Biotechnology and Genomics (CBGP) (SEV-2016-0672). AIFS acknowledges funding received by CONACYT in the form of a Graduate Studies Scholarship. GTdS and MMA gratefully acknowledge the Academic Scholarships provided by CONACyT as members of the National System of Researchers (Sistema Nacional de Investigadores). We thank MSc. Regina E. Vargas-Mejía for her technical assistance during SEM sessions. We acknowledge M.Sc. Sara Cristina Pedroza González for their technical assistance during cell culture experiments.

