## Supplementary Material for "Biofabrication of muscle fibers enhanced with plant viral nanoparticles using surface chaotic flows"

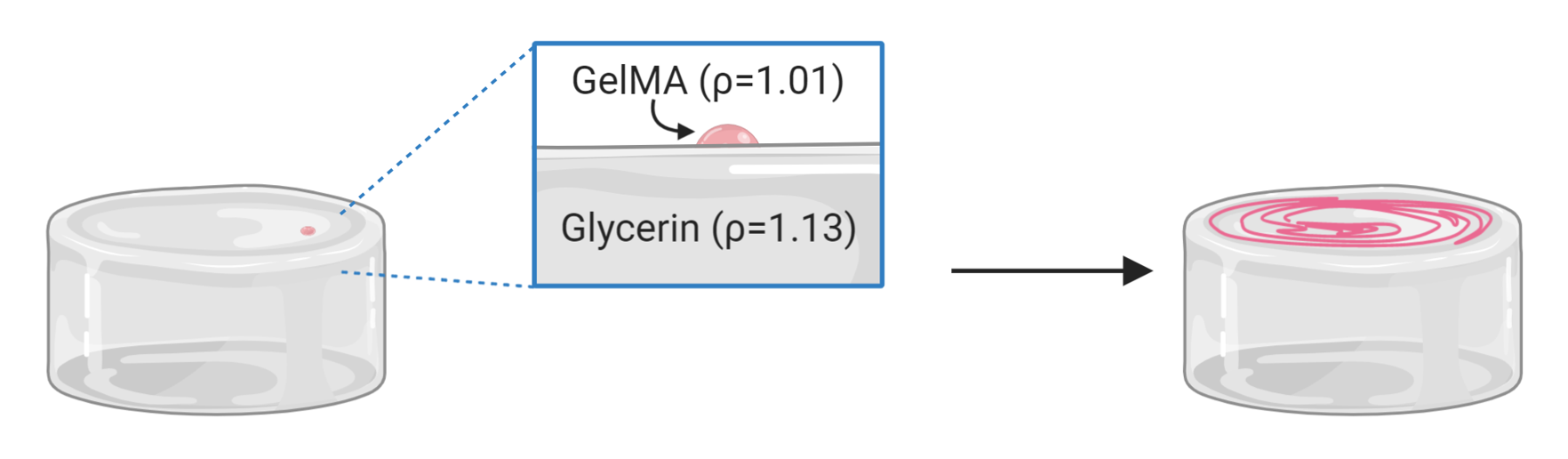

**Figure S1. Difference in densities between working fluids.** The drop of GelMA remains on the surface of the bed of glycerin during the whole process due to their difference in density.

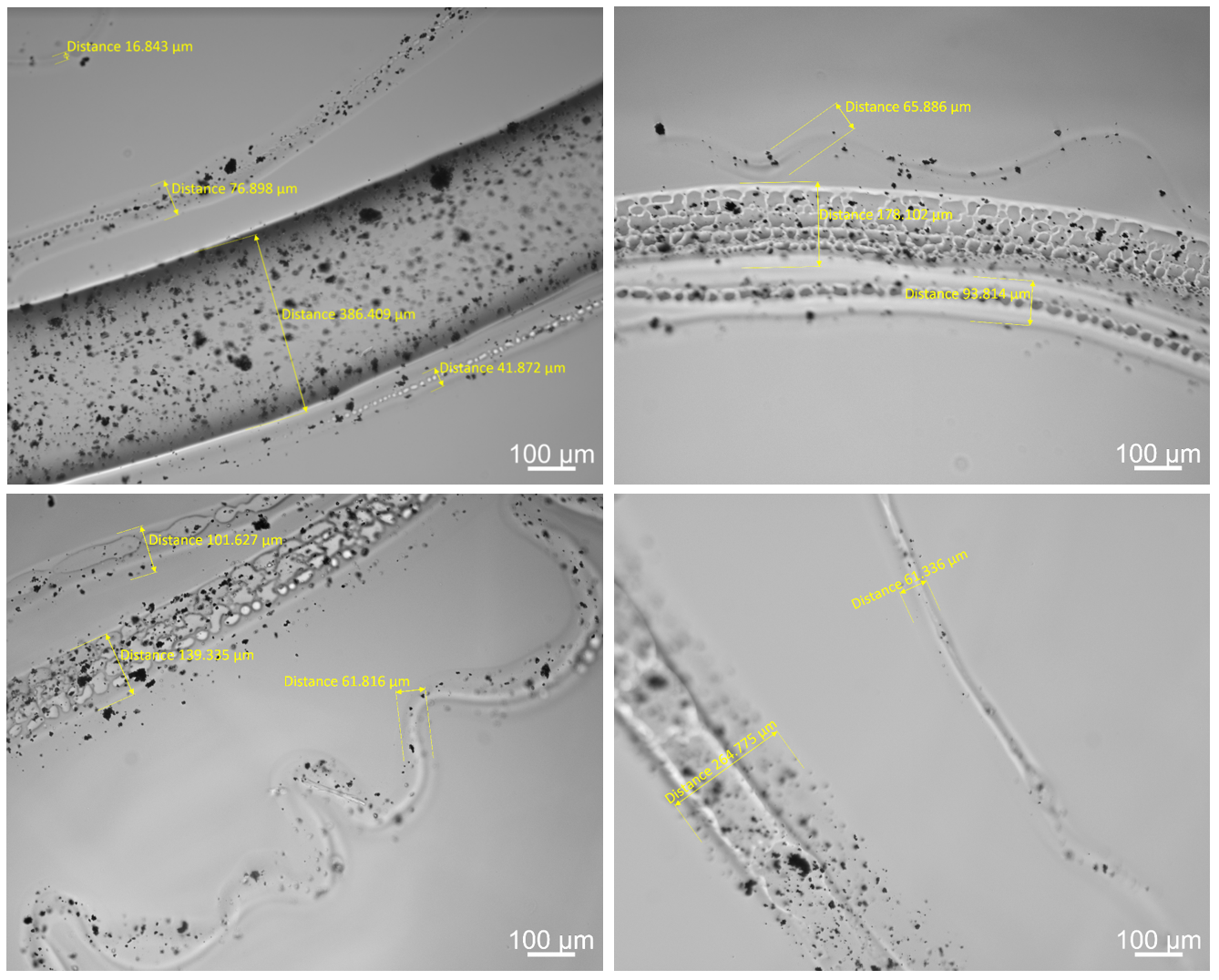

**Figure S2. Width diversity in brightfield micrographs of chaotically printed fibers.** Different field-of-view showing an important diversity of widths in single printed fibers, ranging from 16.8 to 386.4 µm.

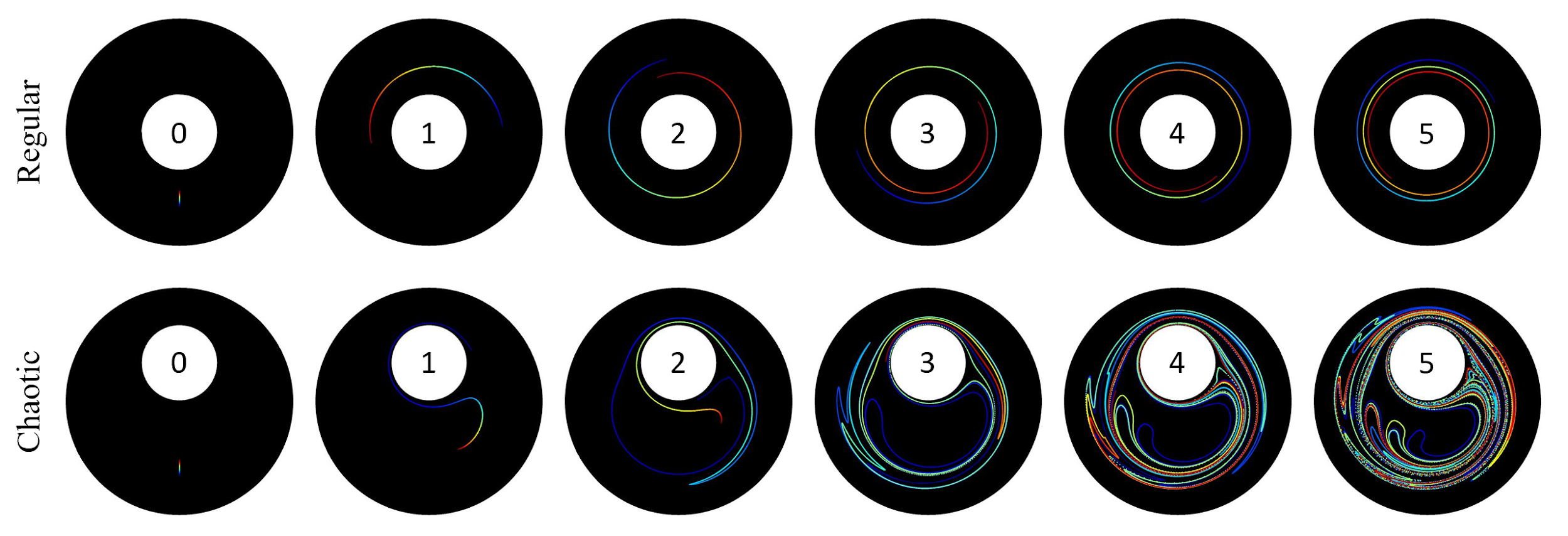

**Figure S3. Progression of ink deformation produced by regular and chaotic flows.** CFD simulations of the deformation of 1 mm lines caused by regular and chaotic flows throughout 5 cycles.

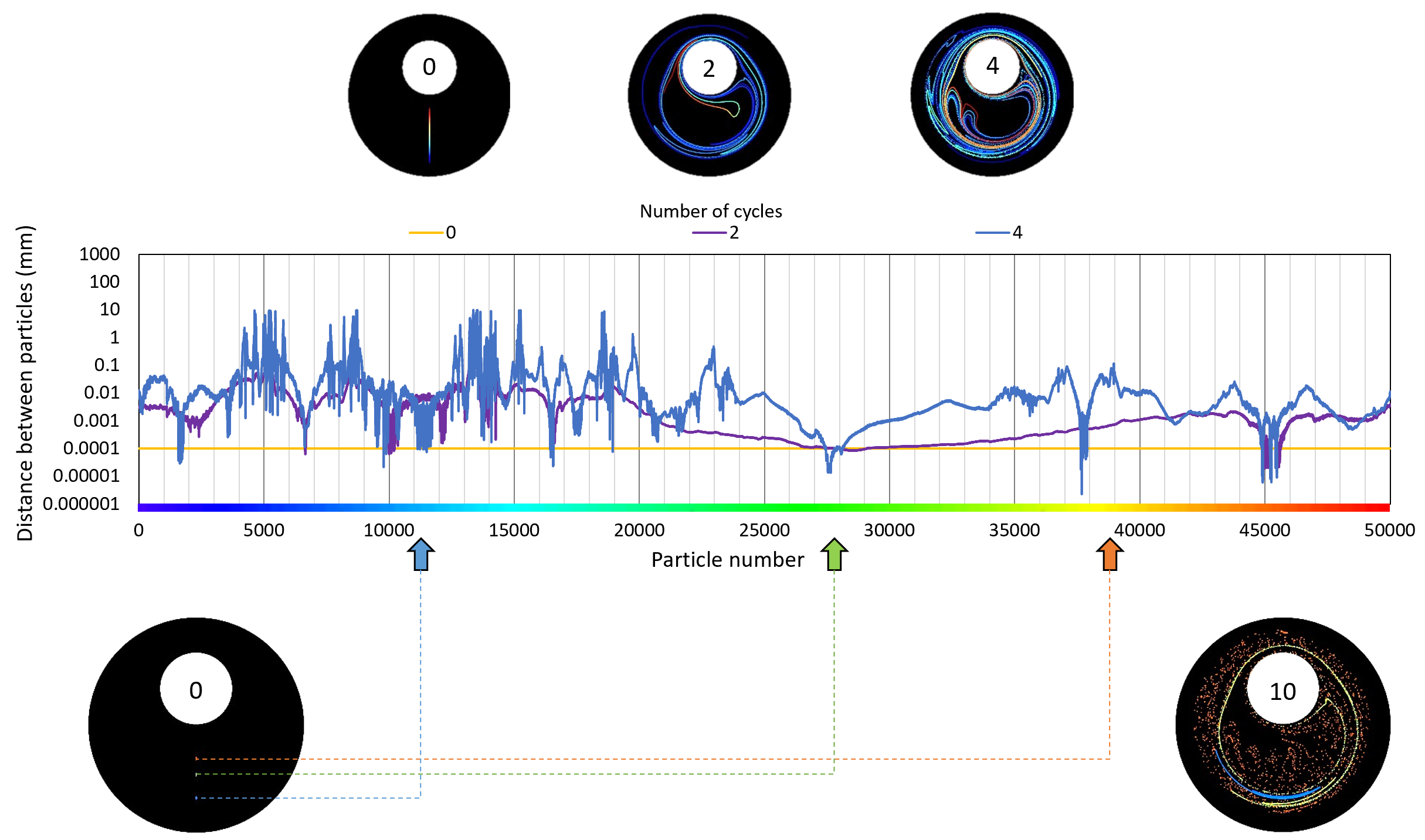

**Figure S4. Effect of the initial ink position on elongation.** Distances between neighboring particles along the centerline at initial condition (yellow), and after the second (purple) and fourth (blue) cycles for the protocol [270°,810°]. Initially all particles are located 0.1 µm apart from each other. Three different initial drop positions in blue, green, and orange, are highlighted, as well as their evolution after 10 cycles.

**Table S1. Experimental fiber lengths.** Experimental measurements from lengths of fibers fabricated through regular and chaotic flows at different numbers of cycles.

| **Cycle *n*** | **Chaotic length (µm)** | **Regular length (µm)** |
| --- | --- | --- |
| 0.0 | 2,684.60 ± 12.14 | 2,684.60 ± 12.14 |
| 0.5 | 13,564.55 ± 447.86 | 14,519.65 ± 871.58 |
| 1.0 | 46,632.92 ± 4,426.09 | 38,253.77 ± 1,116.01 |
| 1.5 | 102,507.78 ± 1,786.41 | 58,385.91 ± 1,528.96 |
| 2.0 | 152,246.01 ± 21,121.50 | 75,972.51 ± 643.49 |

**Table S2. CFD and experimental elongation comparison.** Simulated and experimental elongation coefficients (L/L_0_) of fibers fabricated by chaotic 2D-printing throughout different numbers of cycles.

| **Cycle *n*** | **CFD_C_** | **Experimental_C_** | **CFD_R_** | **Experimental_R_** |
| --- | --- | --- | --- | --- |
| 0.0 | 1.00 | 1.000 ± 0.01 | 1.00 | 1.00 ± 0.01 |
| 0.5 | 3.64 | 5.053 ± 0.17 | 5.45 | 5.41 ± 0.33 |
| 1.0 | 14.41 | 17.371 ± 1.65 | 10.75 | 14.25 ± 0.42 |
| 1.5 | 17.60 | 38.184 ± 0.67 | 16.09 | 21.75 ± 0.57 |
| 2.0 | 65.52 | 56.711 ± 7.87 | 21.42 | 28.30 ± 0.24 |
